# Neighbor cells restrain furrowing during epithelial cytokinesis

**DOI:** 10.1101/2023.06.08.544077

**Authors:** Jennifer Landino, Eileen Misterovich, Shahana Chumki, Ann L. Miller

**Affiliations:** Department of Molecular, Cellular, and Developmental Biology, University of Michigan, Ann Arbor; Cellular and Molecular Biology Graduate Program, University of Michigan, Ann Arbor

**Keywords:** Cytokinesis, Epithelium, Rho GTPase, Actin, Myosin II, *Xenopus*, Optogenetics

## Abstract

Cytokinesis challenges epithelial tissue homeostasis by generating forces that pull on neighboring cells *via* cell-cell junctions. Previous work has shown that junction reinforcement at the furrow in *Xenopus laevis* epithelia regulates the speed of furrowing^1^. This suggests the cytokinetic array that drives cell division is subject to resistive forces from epithelial neighbor cells. We show here that contractility factors accumulate in neighboring cells near the furrow during cytokinesis. Additionally, increasing neighbor cell stiffness, *via* ɑ-actinin overexpression, or contractility, through optogenetic Rho activation in one neighbor cell, slows or asymmetrically pauses furrowing, respectively. Notably, optogenetic stimulation of neighbor cell contractility on both sides of the furrow induces cytokinetic failure and binucleation. We conclude that forces from the cytokinetic array in the dividing cell are carefully balanced with restraining forces generated by neighbor cells, and neighbor cell mechanics regulate the speed and success of cytokinesis.

**Highlights:**

- Neighboring cells assemble actomyosin arrays adjacent to the cytokinetic furrow
- Overexpression of an F-actin cross-linker in neighbor cells slows furrowing
- Optogenetic activation of contractility in one neighbor pauses furrow ingression
- Hyper-contractility in both neighbors restrains furrowing & cells fail cytokinesis

## Introduction

Epithelia act as a biological barrier, defining the external and internal environment of an organism. These tissues consist of polarized epithelial cells that are connected *via* cell-cell junctions, which play an essential role in maintaining homeostasis. In vertebrates, the apical junctional complex includes tight junctions (TJs), responsible for epithelial barrier function, and adherens junctions (AJs), which facilitate cell adhesion and mechanical force transmission between neighboring cells^2, 3^. Apical cell-cell junctions connect to an apical actomyosin array, creating a network of mechanosensitive cells in the epithelium^4^. Epithelia regularly experience mechanical stresses, including those that occur during cell division^5^. Cytokinesis, which physically divides the cell into two daughters, poses a fundamental challenge to epithelial homeostasis. The formation and closure of a cytokinetic furrow, driven by actomyosin-generated forces, induces significant cell shape changes in both the dividing cell and its neighbors^1, 6^. Epithelial cells respond by remodeling and reinforcing their apical cell-cell junctions, ensuring cell adhesion and the maintenance of epithelial barrier function during cell division^1^.

Our current understanding of epithelial cell division has been informed by studies across model systems and tissue types. In *Drosophila*, local adhesion disengagement at the cytokinetic furrow, coupled to contractile actomyosin forces in the neighbor cells leads to the formation of a new junctional interface between daughters^7–9^. Additionally, actomyosin flows in neighbor cells regulate furrow closure, supporting a model where neighbors mechanically regulate cell division^8, 9^. In vertebrate models, dividing cells maintain adhesion at the furrow, pulling neighbor cells inwards, thereby resulting in a four-way junction that can remodel to produce various daughter cell geometries^1, 6^. In *Xenopus*, AJs are specifically reinforced at the cytokinetic furrow, as evidenced by reduced turnover of AJ proteins at the furrow relative to polar regions or interphase cells^1^. This reinforcement relies on the recruitment of the mechanosensitive protein Vinculin^1, 10^, indicating that the cytokinetic contractile array generates high tension within junctional complexes at the furrow. Disrupting Vinculin recruitment accelerates furrow ingression^1^, undermining tension-mediated junction reinforcement and suggesting a direct relationship between junctional forces and the rate of furrow ingression. In cultured epithelial monolayers, a related mechanosensitive recruitment of Vinculin to neighbor cells maintains epithelial integrity during mitotic rounding^11^. More broadly, changes in epithelial tissue tension and stiffness affect cell division orientation and proliferation rates in vertebrate epithelia^12–15^. However, the direct mechanical regulation of furrow closure by neighbor cells in intact vertebrate epithelium remains untested.

Epithelial mechanics are orchestrated, in part, by the actin cytoskeleton, which provides structural support for the apical junctional complex and the apical cell cortex. The small GTPase Rho plays a key role in regulating actomyosin contractility. Active Rho drives assembly of filamentous actin (F-actin) and Myosin II bundles and is enriched at apical cell-cell junctions. Active Rho also promotes assembly and subsequent contraction of the cytokinetic array at the cell equator during anaphase^16–22^. In this study, we examine how the actomyosin cytoskeleton in epithelial neighbor cells controls cytokinesis. Using *Xenopus* embryos as a model system and a combination of live imaging, mosaic expression, and genetic and optogenetic approaches to manipulate the actomyosin array in neighbor cells, we find that Rho-mediated contractility in neighbor cells restrains furrow closure. This demonstrates that non-dividing neighbor cells actively modulate the speed and success of cytokinesis in the vertebrate epithelium.

## Results

### Contractility factors accumulate near the cytokinetic furrow in epithelial neighbor cells

To investigate neighbor cells’ potential to generate a restraining force during cytokinesis, we assessed the localization of contractility factors near the furrow within neighbor cells. Using a mosaic expression approach, we generated an epithelium with mixed cell populations expressing probes for active Rho (rGBD), Myosin II (Sf9 intrabody), and F-actin (LifeAct)^23–25^ as well as non-expressing cells, in early-gastrula stage *X. laevis* embryos. To mitigate the strong accumulation of the contractility factors at the cytokinetic array that overshadow the neighbor cell-specific signal, we selected cell division events where the dividing cell lacked probe expression, while one or more neighboring cells expressed the probes. Using live, confocal microscopy, we identified transient, local increases in active Rho, Myosin II, and F-actin at and near the neighbor cell-cell junction adjacent to the cytokinetic furrow (Figure 1A, 1C, Movies S1 and S2, Figure S1A-C). Kymographs tracking furrow ingression over time revealed that the accumulation of active Rho in neighbors correlated temporally to Myosin II and F-actin accumulation (Figure 1B, 1D). We quantified the signal intensity of active Rho, Myosin II, and F-actin at and near the neighbor cell junction during furrowing (Figure S1F) in cell division events where the time to complete cytokinesis was similar (Figure S1D-E). The intensity of each contractility factor increased in neighbor cells near the furrow during cytokinesis (Figure 1E-F). These findings indicate that neighboring cells in vertebrate epithelium activate or recruit contractility factors at junctions adjacent to the cytokinetic furrow, potentially generating restraining forces *via* Rho and actomyosin in response to forces generated by the cytokinetic contractile ring.

**Figure 1:**
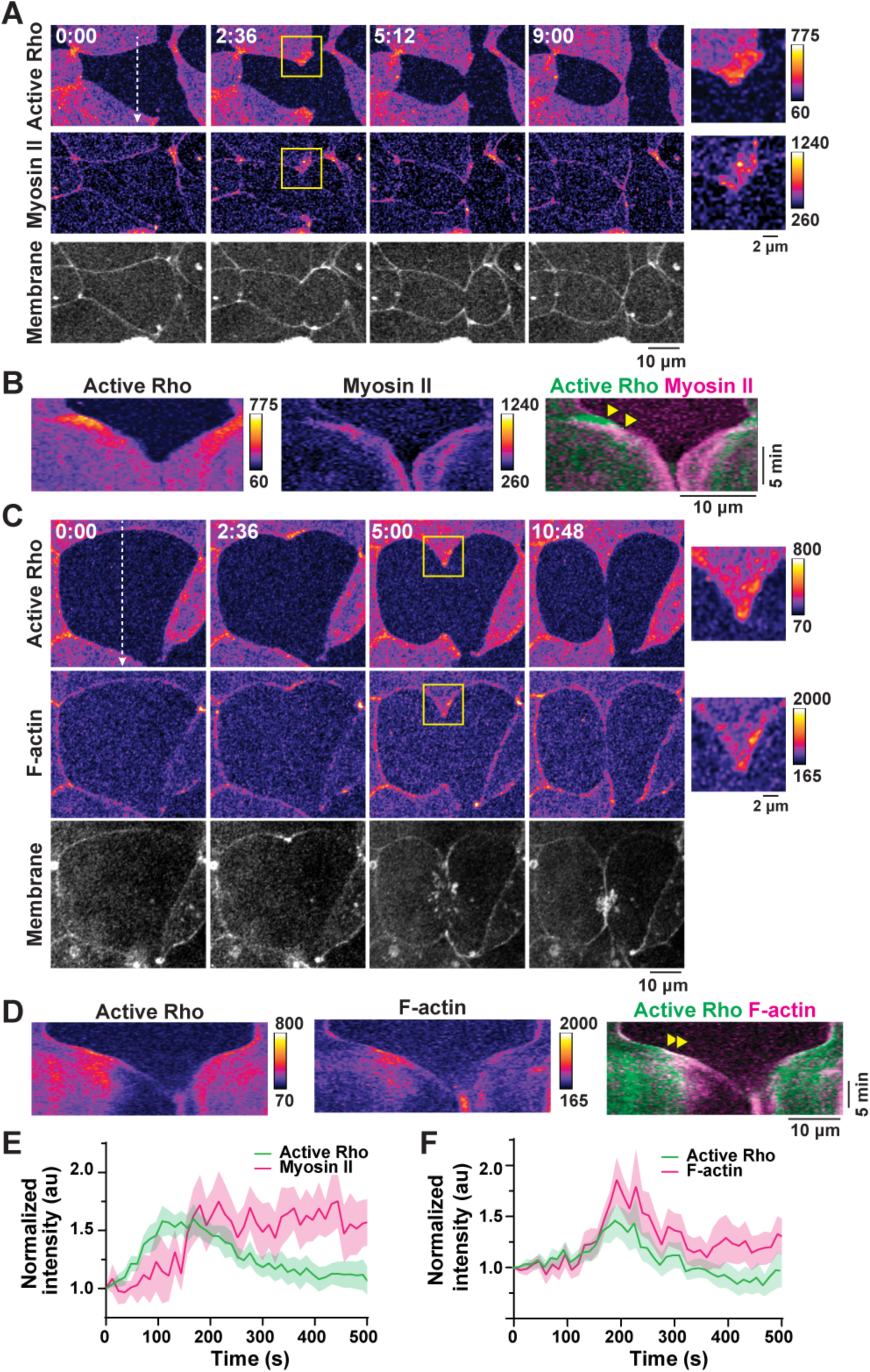
Contractility factors accumulate at and near neighbor cell junctions adjacent to the cytokinetic furrow. A) Micrographs from time-lapse confocal imaging of cytokinesis in an early gastrula-stage *Xenopus laevis* embryo expressing probes for active Rho (GFP-rGBD) and Myosin II (Sf9-mCherry) and stained with a membrane dye (CellMask Deep Red). Yellow boxes correspond to enlarged images shown on the right. Time shown in minutes:seconds from the start of furrow ingression. B) Kymographs of active Rho and Myosin II in neighbors over time generated from the dashed line shown in (A). Yellow arrowheads indicate peak active Rho and Myosin II intensity. C) Micrographs from time-lapse confocal imaging of cytokinesis in an embryo expressing probes for active Rho (GFP-rGBD) and F-actin (LifeAct-RFP) and stained with a membrane dye (CellMask Deep Red). Yellow boxes correspond to enlarged images shown on the right. Time shown in minutes:seconds from the start of furrow ingression. D) Kymographs of active Rho and F-actin in neighbors over time generated from the dashed line shown in (C). Yellow arrowheads indicate peak active Rho and F-actin intensity. E-F) Quantification of active Rho and Myosin II intensity (E) or active Rho and F-actin intensity (F) in neighbor cells where time = 0 is the start of cytokinesis. Intensity was normalized to the membrane signal and the start of ingression (see Figure S1F). Mean (solid line) ± standard error of the mean (SEM, shading). For (E): n = 16 junctions, 14 cells, 5 embryos. For (F): n = 12 junctions, 10 cells, 9 embryos.

### Increasing tissue stiffness in neighbor cells slows cytokinetic furrow ingression

We then asked how manipulation of the actomyosin array within neighbor cells impacts cytokinetic furrow ingression. We hypothesized that hyper-cross-linked F-actin in neighbors would further restrain the forces generated by the cytokinetic contractile ring, thereby slowing furrow closure. To test this, we mosaically overexpressed GFP-tagged ɑ-actinin, a F-actin cross-linker^26–, 28^, which was previously shown to increase tissue stiffness in developing *Xenopus* embryos^29^. We expressed GFP-ɑ-actinin at two levels: 1) a “low ɑ-actinin” that is sufficient for visualization and 2) a “high ɑ-actinin” that is approximately a 4-fold increase in expression. (Figure 2A, Movie S3).

**Figure 2:**
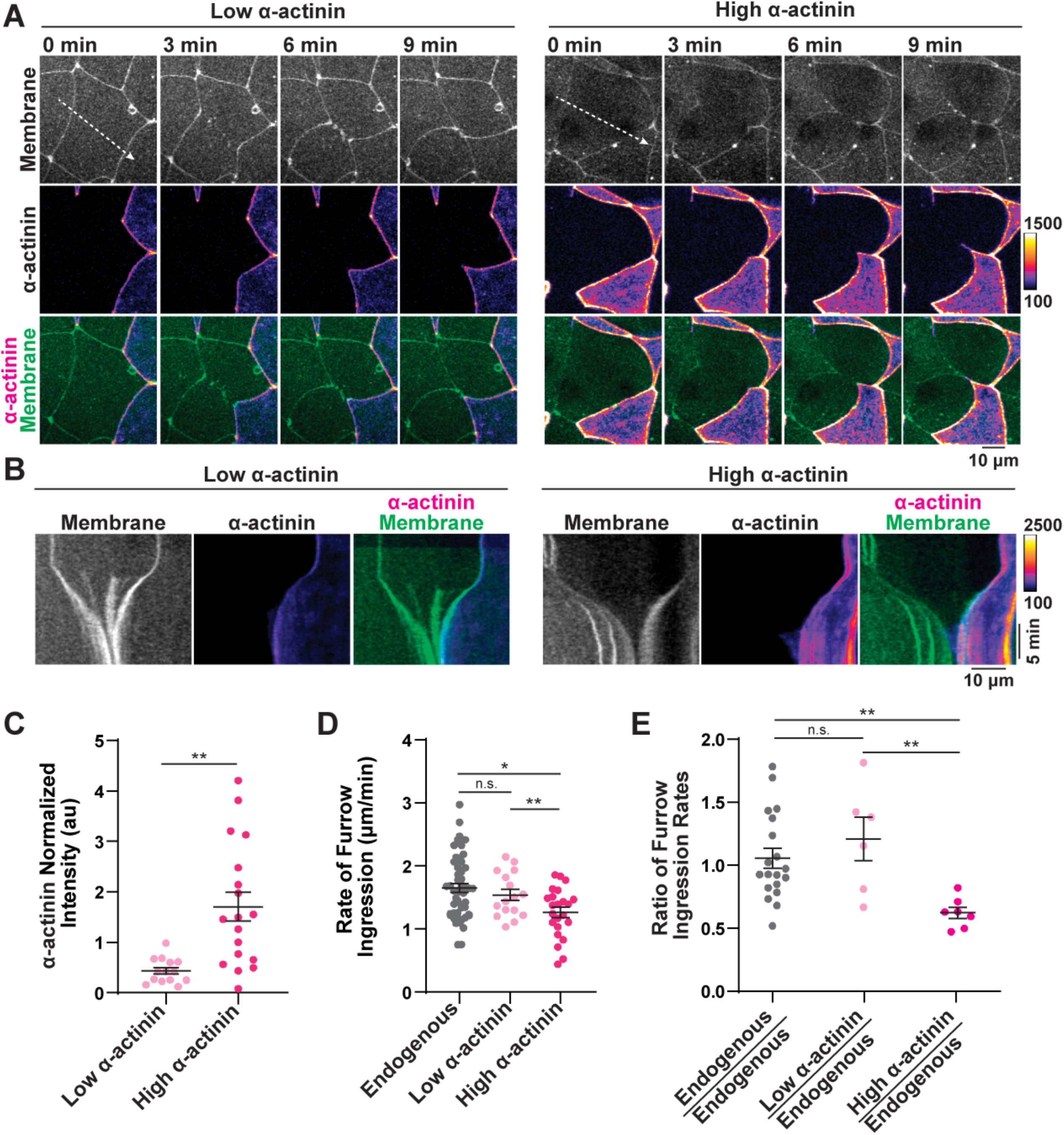
Overexpression of ɑ-actinin in neighbor cells slows cytokinetic furrow ingression. A) Micrographs from time-lapse confocal imaging of cytokinesis with neighbors overexpressing low or high levels of GFP-ɑ-actinin and stained with a membrane dye (CellMask Deep Red). B) Kymographs of membrane and ɑ-actinin signal during furrowing generated from the dashed lines shown in (A). C) Quantification of ɑ-actinin signal intensity normalized to membrane in neighbor cells used in the analysis of rate of furrow ingression. Mean ± SEM. For low overexpression: n = 14 cells, 5 embryos. For high overexpression: n = 18 cells, 7 embryos. ** p ≤ 0.005. D) Quantification of the rate of furrow ingression for junctions with neighbors expressing endogenous, low overexpression, or high overexpression of ɑ-actinin. Mean ± SEM. For junctions with endogenous neighbors: n = 50, 32, 19 (junctions, dividing cells, embryos), for low overexpression: n = 16, 11, 5, for high overexpression: n = 23, 17, 8. * p ≤ 0.05 and ** p ≤ 0.005. E) Ratio of furrow ingression rate for cells with two neighbors with endogenous levels of ɑ-actinin, one endogenous and one low overexpression neighbor, and one endogenous and one high overexpression neighbor. Mean ± SEM. For two endogenous neighbors: n = 19 cells, 10 embryos, for one low overexpression/one endogenous neighbor: n = 6 cells, 3 embryos, for one high overexpression/one endogenous neighbor: n = 7 cells, 6 embryos. ** p ≤ 0.005.

The normalized whole-cell signal intensity of ɑ-actinin in neighbor cells was 0.44 ± 0.07 (low ɑ-actinin) and 1.71 ± 0.29 (high ɑ-actinin, mean ± SEM, Figure 2C). We tracked cell division events with one or more neighbor cells expressing GFP-ɑ-actinin and independently quantified furrow ingression rate for each cell-cell junction based on the neighbor cell’s ɑ-actinin level (endogenous, low, or high ɑ-actinin, Figure S2A) using kymographs (Figure 2B). The furrow ingression rate for junctions with endogenous ɑ-actinin neighbors was not significantly different from those with low ɑ-actinin neighbors (1.65 ± 0.07 µm/min for endogenous neighbors, 1.54 ± 0.35 µm/min for low ɑ-actinin neighbors, mean ± SEM, Figure 2D). However, junctions with high ɑ-actinin neighbors had a significantly slower ingression rate compared to endogenous or low ɑ-actinin neighbors at 1.27 ± 0.39 µm/min (Figure 2D). We also compared the ratio of furrow ingression rates for cell division events with two different neighbors (one endogenous control and one ɑ-actinin overexpression neighbor, Figure S2A). If restraining forces generated from the neighbor cells are equal, we expect the ratio of furrow ingression rates to be close to 1. This was observed for dividing cells with two endogenous neighbors (1.06 ± 0.08, mean ± SEM, Figure 2E). Dividing cells with one endogenous and one low ɑ-actinin neighbor showed a slightly increased ratio, but not significantly different from cells with two endogenous neighbors (1.35 ± 0.14). However, dividing cells with one endogenous and one high ɑ-actinin neighbor exhibited a significantly reduced ratio (0.62 ± 0.12), indicating that highly cross-linked F-actin in a single neighbor generates enough restraining force to induce asymmetric furrow closure. Taken together, these results demonstrate that increasing actomyosin stiffness in epithelial neighbor cells slows the rate of cytokinetic furrow ingression.

### Optogenetic activation of Rho-mediated contractility in one neighbor cell pauses furrow ingression

We next investigated the impact of acute changes in neighbor cell contractility on cytokinetic furrow ingression, using the TULIP optogenetic system^30–33^ (Figure S3A). By locally activating Rho in neighbor cells on one side of the furrow during cytokinesis, we induced a real-time, rapid change in neighbor cell contractile output. Using a probe for active Rho as a junctional marker^24^, we first verified that the TULIP optogenetic system increased Rho activation upon light stimulation (Figure S3B). We measured changes in active Rho signal over time in three areas: 1) a region of light stimulation on one side of the furrow (stimulated neighbor), 2) an unstimulated region on the opposite side of the dividing cell (unstimulated neighbor), and 3) a distant region in the field of view (control region, Figure S3B). Within the region of the stimulated neighbor, active Rho signal intensity peaked approximately 269 seconds after the start of stimulation with a maximum 1.31 ± 0.03-fold change in intensity (mean ± SEM) relative to the whole field active Rho intensity. In the region of the unstimulated neighbor, we observed a small increase in the active Rho signal, which peaked 278 seconds after the start of stimulation (maximum 1.18 ± 0.06-fold change). Notably, active Rho signal increase was significantly higher for stimulated neighbors compared to unstimulated neighbors at 148-269 seconds after the start of stimulation. We speculate that the modest increase in active Rho in the unstimulated neighbor is due to a tension transmission response, causing Rho accumulation at junctions near the site of local stimulation as shown previously^33^. The control region showed no significant increase in active Rho (1.08 ± 0.04-fold higher than the whole field active Rho intensity 278 seconds after the start of stimulation, Figure S3B), confirming the local activation of Rho in optogenetically stimulated neighbor cells.

To test whether acute changes in neighbor cell contractility impact cytokinesis, we allowed furrow ingression to begin, then stimulated neighbors on a single side of the furrow to activate Rho-mediated contractility for approximately five minutes. We stopped stimulation while monitoring ingression by live imaging (Figure 3A, Movie S4). We observed a pause in furrowing on the stimulated side of the furrow, while the unstimulated side continued to ingress (Figure 3B). Despite the temporary pause in furrowing caused by neighbor cell stimulation, all cells successfully completed cytokinesis after stimulation ended. Using kymographs to track the furrow’s position over time, we quantified ingression rate for each side of the furrow before, during and after optogenetic stimulation (Figure S3C). The rate of furrowing significantly decreased during neighbor cell stimulation, then returned to normal after stimulation ended (0.56 ± 0.08 µm/min before, 0.08 ± 0.17 µm/min during, 0.85 ± 0.40 µm/min after stimulation, mean ± SEM, Figure 3C). There was no significant change in the rate of furrow ingression on the unstimulated side of the dividing cell before (0.94 ± 0.26 µm/min), during (0.79 ± 0.23 µm/min), or after stimulation ((0.77 ± 0.17 µm/min). We also quantified the total rate of furrowing for junctions in dividing cells with light-stimulated neighbors and control cells that did not experience neighbor stimulation (Figure S3C). Both the stimulated (0.53 ± 0.09 µm/min) and unstimulated (0.76 ± 0.12 µm/min) sides of the furrow had a significantly decreased furrow ingression rate compared to control cells without stimulated neighbors (1.29 ± 0.12 µm/min, mean ± SEM, Figure 3D). Therefore, an acute increase in contractility of a single neighbor cell can slow the overall rate of furrow ingression relative to cells without light-stimulated neighbors, demonstrating the impact of neighbor cell hypercontractility on furrow dynamics. Our findings show that induction of Rho-mediated contractility in one neighbor cell can asymmetrically pause cytokinetic furrowing, extending the time required to complete cytokinesis. Importantly, a brief pause in furrow ingression does not inhibit successful cell division.

**Figure 3:**
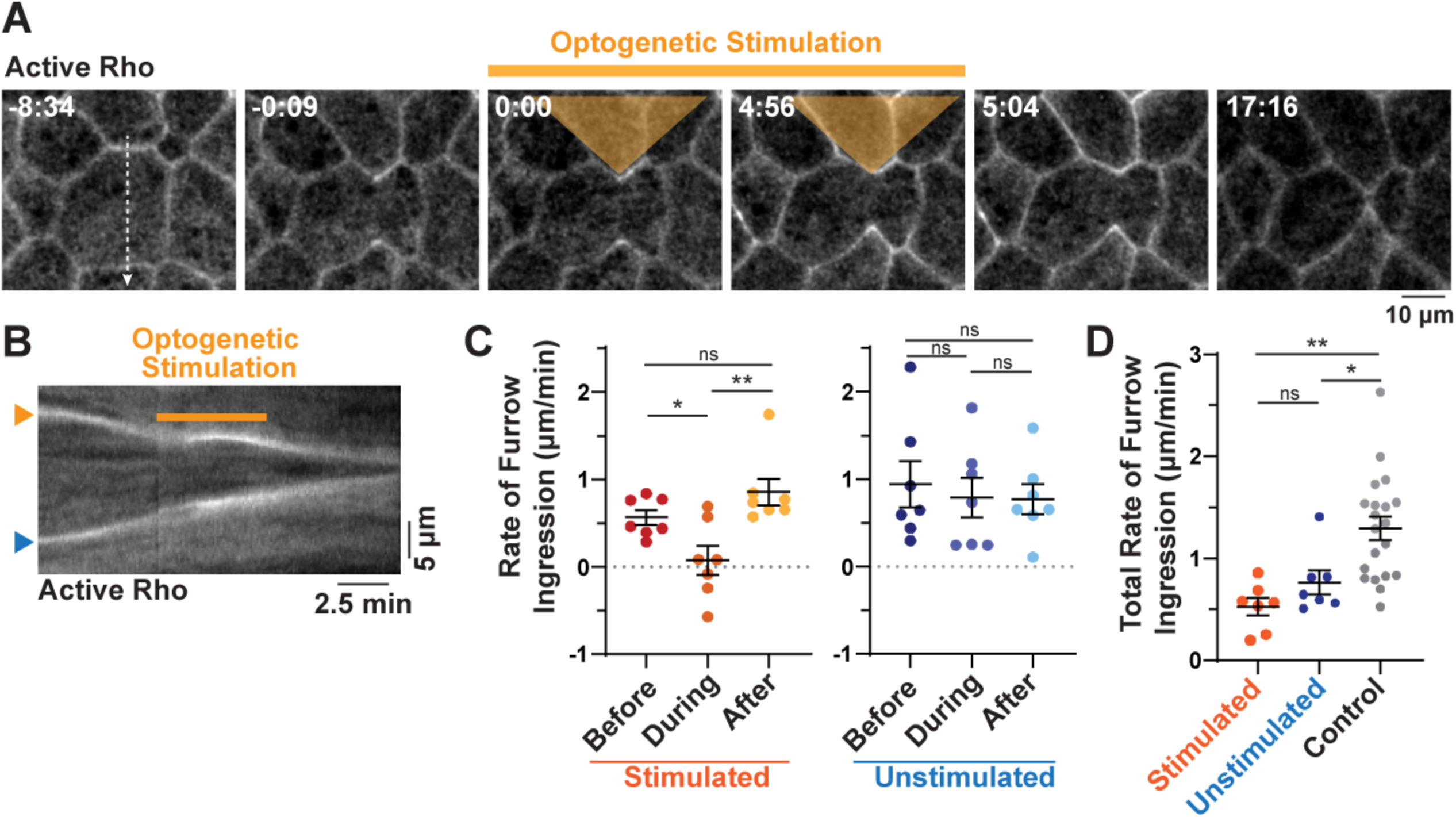
Optogenetic activation of contractility in one neighbor cells stalls furrowing. A) Micrographs from time-lapse confocal imaging of cytokinesis with optogenetic stimulation of Rho- mediated actomyosin contractility in neighbors on one side of the furrow. Orange triangle and orange line indicate the region and duration of stimulation, respectively. Time shown in minutes:seconds. B) Kymograph of furrow ingression generated from the dashed line shown in (A). Orange arrowhead indicates side of furrow with a stimulated neighbor, blue arrowhead indicates side of furrow with unstimulated neighbor. Orange line indicates the duration of stimulation. C) Quantification of the rate of furrow ingression before, during, and after neighbor cells were optogenetically stimulated. Junctions neighboring the stimulated cells are shown in warm colors, while corresponding junctions with unstimulated neighbors are shown in cool colors. Mean ± SEM. n = 7 junctions, 7 cells, 5 embryos. * p ≤ 0.05 and ** p ≤ 0.005. D) Total rate of furrow ingression for junctions from cells with stimulated (orange) or unstimulated (blue) neighbors, as shown in (C), or for junctions in control cells selected from a region in the field of view that was well separated from the region of optogenetic stimulation. Mean ± SEM. For stimulated and unstimulated junctions: n = 7 junctions, 7 cells, 5 embryos, for control cells: n = 20 junctions, 10 cells, 6 embryos. * p ≤ 0.05 and ** p ≤ 0.005.

### Optogenetic activation of Rho-mediated contractility in both neighbor cells induces cytokinetic failure

As cells exit mitosis there is a brief window where the cortex is competent to contract^34^. We hypothesized that stalling ingression on both sides of the furrow during this window would result in failed ingression and binucleation. To test this, we optogenetically activated Rho-mediated contractility in neighbor cells on both sides of the furrow shortly after ingression began and monitored mitotic exit using a chromatin marker (H2B) (Figure 4A). We found that stimulation of both neighbors caused the cytokinetic furrow to stall or regress in most cases, and the furrow maintained this position until the end of the 10-minute stimulation period (Figure 4A-B). For cells that stalled or regressed, we measured the furrow width at three time points: 1) during anaphase, 2) at maximum ingression, and 3) after 10 minutes of stimulation (Figure 4B). We found that the cytokinetic furrow ingressed to 74.12 ± 2.79% (mean ± SEM, Figure 4C) of the initial cell width during anaphase. After stimulation, the cell width at the equator was 85.0 ± 5.05% of the width in anaphase (Figure 4C), indicating that stimulation of both neighbor cells was sufficient to block completion of cytokinetic furrowing. After 10-minutes of stimulation, cells were unable to resume ingression, and chromosomes decondensed, indicating entry into the next cell cycle. We quantified the cytokinetic success rate for cells with two stimulated neighbors by determining the percentage of cells that completed cytokinesis with two distinct daughter cells. Cells with two optogenetically stimulated neighbors completed cytokinesis 23.33 ± 14.53% of the time (mean ± SEM, Figure 4D), while dividing cells without stimulated neighbors in the same field of view had a success rate of 82.78 ± 8.75 (Figure 4D). Cells expressing the optogenetic constructs but undergoing division in non-stimulated epithelial tissue had a cytokinesis success rate of 78.33 ± 11.67% (Figure 4D). This indicates that acute changes in contractility of both neighbor cells resists cytokinetic furrow ingression, leading to frequent cell division failure and binucleation.

**Figure 4:**
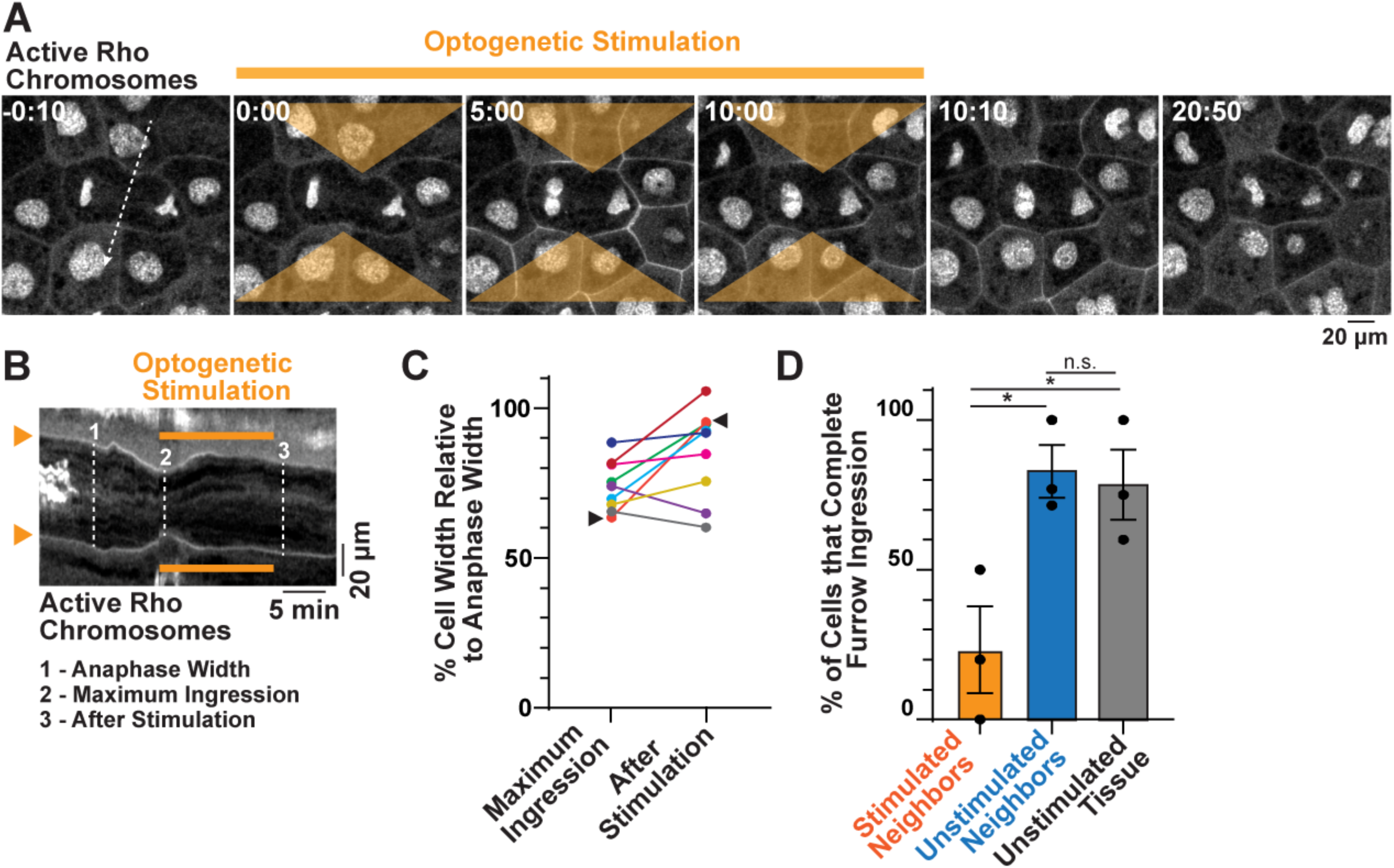
Optogenetic activation of contractility in both neighbors causes cytokinetic failure. A) Micrographs from time-lapse confocal imaging of cytokinesis with optogenetic stimulation of Rho-mediated actomyosin contractility in neighbors on both sides of the furrow. Orange triangles and orange line indicate the regions and duration of stimulation, respectively. Time shown in minutes:seconds. B) Kymograph of furrow ingression generated from the dashed line shown in (A). Orange arrowheads indicate both sides of the furrow with a stimulated neighbor. Orange line indicates the duration of stimulation. Dashed lines indicate example time points used to measure cell width at Anaphase (1), Maximum Ingression (2), and After Stimulation (3). C) Quantification of cell width at the equator relative to the initial anaphase width. Arrowheads indicate the maximum ingression width and width after stimulation for the cell shown in (B). D) Quantification of the success of cytokinesis for dividing cells with stimulated neighbors, dividing cells in the same field of view with unstimulated neighbors, and cells expressing the optogenetic system but dividing in tissue that did not experience light stimulation. Mean ± SEM. For stimulated neighbors: n = 12 cells, 12 embryos, for unstimulated neighbors: n = 28 cells, 10 embryos, and for unstimulated tissue: n = 26 cells, 8 embryos, * p ≤ 0.05.

## Discussion

Forces generated by dividing cells in epithelia underlie tissue architecture, homeostasis, and cell polarity and division orientation^1, 13, 35–37^. Epithelial cytokinesis requires dynamic remodeling of cell-cell junctions and neighbor cell geometry to accommodate the formation and closure of the cytokinetic furrow. How neighbor cells respond to forces transmitted across cell-cell junctions at the furrow and whether neighbors mechanically regulate cytokinesis in the vertebrate epithelium has not been investigated. Here, we show that neighbor cells respond to force generated by the cytokinetic contractile ring *via* local accumulation of contractility factors adjacent to the furrow, with spatiotemporal patterning that is consistent with a Rho-mediated actomyosin response. Modulation of the neighbor cell cytoskeletal array, either through genetic overexpression of the crosslinking protein ɑ-actinin or the optogenetic activation of Rho, slows or stalls furrowing. Prolonged stalling of furrow ingression due to acute increases in neighbor cell contractility on both sides of the furrow overrides the force generated by the cytokinetic contractile array, resulting in cell division failure.

Actomyosin accumulation in neighbor cells during furrowing in the *Xenopus* epithelium mirrors the *Drosophila* response where epithelial neighbors locally accumulate Myosin II near the furrow^8, 9^. Despite differences in neighbor cell geometry, cell interface rearrangements, and junctional protein dynamics at the furrow between these model systems^1, 37^, this suggests that multicellular regulation of epithelial cytokinesis *via* neighbor cell contractility is evolutionarily conserved. However, the function of neighbor cell contractility may be different between tissues, as Myosin II accumulation in *Drosophila* neighbors is necessary to form a new daughter-daughter interface while cytokinesis in vertebrate models most often results in a new four-way junction^1, 6, 8, 9^. Our data suggests that neighbor cell contractility in *Xenopus* regulates the speed of furrowing. The upstream signal activating actomyosin assembly in neighbor cells may also differ. Recent work in *Drosophila* has shown that a local decrease in E-cadherin at the furrow serves as the mechanical signal for neighbor cell RhoGEF activation and actomyosin assembly^8, 38^. In *Xenopus*, E-cadherin is reinforced at the furrow, not reduced^1^, indicating an alternative mechanism of activation for neighbor cell contractility. Identifying the molecules that activate Rho in neighbor cells in response to forces generated by the cytokinetic contractile ring and transmitted across cell-cell junctions to neighbor cells will be an important area for future studies. Likely candidates include mechanosensitive RhoGEFs that localize to apical cell-cell junctions.

Epithelial cytokinesis is fundamental for multicellular organism development and homeostasis^39, 40^. Disruption of well-regulated cell proliferation leads to solid tumor formation, and failure to successfully complete cytokinesis results in aneuploidy, a hallmark of cancer cells^41–43^. We have identified a novel mechanism regulating epithelial cytokinesis: modulation of the contractile array in neighbor cells controls the speed and success of furrow ingression. Recent work has shown that increased tissue strain in oncogenic epithelia alters cell division rates in wild-type neighboring cells^13^, suggesting that neighbor cell mechanics play an important role in homeostasis. Our work demonstrates how manipulating the local tissue environment can impact the outcome of cytokinesis, even when the dividing cell itself is not directly perturbed. Understanding the relationship between cytokinetic forces and neighbor cell actomyosin arrays is essential to define the mechanisms underlying epithelial cytokinesis in development and disease.

## Materials and Methods

### Key Resources Table

| REAGENT or RESOURCE | SOURCE | IDENTIFIER |
| --- | --- | --- |
| Experimental models: Organisms/strains |  |  |
| *Xenopus laevis* (female), oocyte positive, pigmented | Xenopus 1 | N/A |
| *Xenopus laevis* (female), oocyte positive, albino | Xenopus 1 | N/A |
| *Xenopus laevis* (male), mature, pigmented | Xenopus 1 | N/A |
| Recombinant DNA |  |  |
| pCS2+/GFP-rGBD | 23 | N/A |
| pCS2+/mCherry-2XrGBD | 44 | N/A |
| pCS2+/Sf9-mCherry | This paper | N/A |
| pCS2+/Sf9-mNeon | 45 | N/A |
| pCS2+/LifeAct-RFP | 1 | N/A |
| pCS2+/LifeAct-GFP | 1 | N/A |
| pCS2+/GFP-α-actinin | Gift from Lance Davidson, University of Pittsburgh<sup>29,46</sup> | N/A |
| pCS2+/2xPDZ-YFP-LARG(DH) | 33 | N/A |
| pCS2+/Stargazin-GFPsilent-LOVPEP | Gift from Patrick Oakes, Loyola University Chicago<sup>33</sup> | N/A |
| pCS2+/mCherry-H2B | 20 | N/A |
| Software and algorithms |  |  |
| GraphPad Prism |  | N/A |
| Microsoft Excel |  | N/A |
| Fiji (ImageJ) | 47 | N/A |
| Other |  |  |
| CellMask Deep Red Plasma Membrane Stain | ThermoFisher | C10046 |
| mMESSAGE mMACHINE SP6 Transcription kit | ThermoFisher | AM1340 |

### Resource availability Lead contact

Further information and requests for resources and reagents should be directed to and will be fulfilled by the Lead Contact, Ann L. Miller

### Materials availability

Plasmids generated in this study can be obtained from the corresponding author Ann L. Miller

### Experimental model and subject details

Adult *Xenopus laevis* wild type and albino female frogs and wild type male frogs were purchased from Xenopus 1 (Dexter, MI). Female frogs were injected with human chorionic gonadotropin (HCG) to induce them to lay eggs. Male frogs were used for acquisition of testes for sperm preparations.

Frogs were housed in a recirculating tank system (Tecniplast, Milan, Italy), which constantly monitors water quality parameters (temperature, pH, and conductivity) to ensure safe and consistent water quality for an optimal environment for frog health. Daily health and maintenance checks were performed by Animal Care Staff, and frogs were fed a 50/50 mix of Nasco frog brittle + Skretting Protec 2.5 trout diet two times per week.

All studies strictly adhered to the compliance standards of the US Department of Health and Human Services Guide for the Care and Use of Laboratory Animals and were approved by the University of Michigan’s Institutional Animal Care and Use Committee. A board-certified Laboratory Animal Veterinarian oversees our animal facility.

### Method Details

#### *Xenopus laevis* embryos and microinjections

*Xenopus* eggs were collected, fertilized *in vitro*, and de-jellied^19, 48^. De-jellied embryos were kept at room temperature or at 15°C in 0.1X Mark’s modified Ringer’s solution (MMR) containing 10 mM NaCl, 0.2 mM KCl, 0.2 mM CaCl2, 0.1 mM MgCl2, and 0.5 mM Hepes, pH 7.4. Embryos were microinjected in the animal hemisphere with mRNA either twice per cell at the two-cell stage or once per cell at the four-cell stage. For mosaic expression, only one of two, or one of four cells were injected. Injected embryos were allowed to develop to gastrula stage (Nieuwkoop and Faber stage 10.5-12^49^) at 13°C or 15°C. For optogenetic experiments, embryos were kept in the dark after mRNA injection. The amount of mRNA per 5 nl of microinjection volume was as follows: GFP-rGBD, 80 pg for co-expression experiments and 50 pg for single expression experiments; mCherry-2XrGBD, 40 pg; Sf9-mCherry, 140 pg for co-expression experiments, Sf9-mNeon, 85 pg for single expression experiments; LifeAct-RFP, 25 pg for co-expression experiments; LifeAct-GFP, 25 pg for single expression experiments; GFP-ɑ-actinin, 250 pg for low overexpression, 1 ng for high overexpression; GFPsi-LOVpep, 5 pg; prGEF-YFP, 2 pg, mCherry-H2B, 8 pg. To stain the cell membrane, embryos were incubated in CellMask Deep Red Plasma membrane stain (ThermoFisher, Waltham, Massachusetts) at 2 µg/ml in 0.1X MMR for 2 minutes, and then washed for 10 minutes in 0.1X MMR before imaging.

#### mRNA preparation

All plasmid DNAs were linearized with NotI. mRNAs were transcribed *in vitro* using the mMESSAGE mMACHINE SP6 Transcription kit (ThermoFisher), and purified using the RNeasy Mini kit (Qiagen, Venlo, The Netherlands). Transcript size was verified on a 1% agarose gel containing 0.05% bleach and Millenium RNA markers (ThermoFisher).

#### Generation of new DNA constructs

Sf9 was a generous gift from Ed Munro^25^ and was subcloned into the pCS2+/mCherry plasmid as previously described for pCS2+/Sf9-mNeon^45^. Briefly, Sf9 was amplified by PCR using Herc II Fusion polymerase, and cloned into pCS2+/mCherry using BamHI and ClaI restriction enzymes. Sf9 insertion was validated by whole plasmid sequencing (Plasmidsaurus).

### Experimental replicates

Multiple cell division events from multiple embryos were used for all experimental replicates (as indicated in the figure legend). Each experiment was conducted using three or more biological replicates with multiple embryos obtained from individual frogs, except for co-localization of active Rho and Myosin in neighbor cells which was conducted with two experimental replicates.

### Confocal microscopy

Live confocal laser scanning microscopy of cytokinesis in early gastrula-stage *Xenopus laevis* embryos was performed on an inverted Olympus Fluoview 1000 microscope equipped with a 60X supercorrected PLANON 60XOSC objective (numerical aperture [NA] = 1.4; working distance = 0.12 mm) and FV10-ASW software. Embryos were suspended in 0.1X MMR and mounted in a 0.8 mm-thick metal slide with a 10 mm hole in the center, sandwiched between two coverslips adhered with vacuum grease. Generally, imaging of epithelial cytokinesis was performed by collecting three to five z-slices at a step size of 0.5 µm. For imaging cytokinesis during optogenetic stimulation of both neighbors, the sample was imaged by collecting three 1 µm z-slices. For co-imaging experiments, channels were acquired sequentially to minimize bleed through.

### Optogenetic stimulation

Simultaneous live imaging and optogenetic stimulation was performed on the Olympus Fluoview 1000 as described above using the SIM scanner. Imaging of active Rho was performed by scanning three apical z-slices with a step size of 0.5 µm or 1 µm. Simultaneous optogenetic stimulation was performed by creating a triangular region of interest (ROI) that encompassed cells neighboring the cytokinetic furrow. The ROI was stimulated with the SIM scanner using 3% 405 nm laser power for a duration of 1 s with a 20 s interval for a total time of approximately 5 minutes (single neighbor stimulation) or 10 minutes (double neighbor stimulation)^33^.

### Image analysis

Intensity of active Rho, Myosin II, and F-actin in neighbor cells: a circular ROI with a 5 µm diameter was drawn in the neighbor cell encompassing the neighbor cell junction at the cytokinetic furrow and adjacent cytoplasmic area. The active Rho, Myosin II, and F-actin signal intensities were normalized to the membrane signal in the same ROI. All intensity measurements were background corrected using a measurement from a ROI of the same shape and size centered on a non-expressing area of the embryo in the same field of view. The intensity at each time point was normalized to the intensity at the start of cell division (see Figure S1F). For co-expression experiments, each time point in a live imaging series was measured and normalized to the starting time = 0. For single expression experiments, measurements were taken at 0%, 25%, 50%, and 75% completion of furrow ingression. For each percent ingression measurement, a measurement from the frame before, the frame itself, and the frame after were quantified and averaged (example: 25% ingression ± 1 frame) and then normalized to 0% ingression. For non-dividing control cells, we selected cells in the same field of view and measured signal at time points similar to the timeline of cytokinetic progression (0, 250, 500, and 750 seconds), with time = 0 matched to the start of furrowing of the nearby dividing cell. This approach was used to account for drifting in the focal plane that could affect the global signal intensity across the field of view.

Time to complete furrowing: the duration of cytokinesis was determined by calculating the number of frames it took to complete cytokinesis, with the first frame being the time when the furrow began to form and the last frame being the time the two junctions met to form a 4-way intersection.

Intensity of ɑ-actinin in neighbor cells: only neighbor cells that were fully visible within a field of view were used for analysis. The ROI was drawn to encompass the entire neighbor cell and the mean intensity of the ɑ-actinin channel was background corrected using a non-expressing region and normalized to the mean intensity of the membrane channel.

Rate of furrow ingression with control and ɑ-actinin overexpressing neighbor cells: the furrow rate was calculated using kymographs generated using a 5 pixel-wide line across the division plane. The distance moved by the junction was determined by marking the starting point of the furrow moving inward and the end point where the two junctions met and before they became indistinguishable. The change in distance was divided by the change in time to determine the rate of furrow ingression. To determine the ratio of furrow ingression for dividing cells with two different neighbors, the rate of ingression for the ɑ-actinin overexpressing neighbor was divided by the rate of ingression for the control, endogenous expressing neighbor (see Figure S2A).

Rate of furrow ingression during optogenetic stimulation: kymographs were generated from a 5 pixel-wide line across the division plane and the time points corresponding to before, during and after stimulation were marked as start and end points. The distance the junction moved between each of the time points was determined as described above. The rate was calculated as the change in distance divided by the change in time. To calculate the total rate of furrow ingression, a similar approach was used with the starting point as the start of furrow ingression and the end as the time point when the two junctions met and just before they became indistinguishable.

Intensity of active Rho during optogenetic stimulation: triangular ROIs were drawn that correspond to the area and position of optogenetic stimulation (stimulated neighbor). ROIs of the same size were drawn on the unstimulated neighbor cells (mirrored to the orientation of the stimulated neighbor) and in an area well-distanced from the stimulation region (control region). The mean intensity of active Rho in each ROI was measured over time (before, during and after stimulation). The mean intensity was normalized to the whole field mean intensity of active Rho at the corresponding time point, which was measured using a square ROI that encompassed the entire field of view. For embryos with focal drift, only the portion of the embryo that remained in focus for the entire duration of imaging was used for the whole field intensity measurement.

Cell width during optogenetic stimulation of both neighbors: The cell width at the cell equator during anaphase A (before furrow ingression) was measured as the starting cell width. The maximum ingression was measured when the distance across the cell equator was the shortest, and the cell width was measured again at the time point when stimulation ended. The percent cell width was calculated at maximum ingression and after stimulation relative to the anaphase cell width.

Success of furrow ingression: The success or failure of cytokinetic furrow ingression was scored based on the position of the furrow at timepoints ≥ 5 minutes after stimulation or completion of furrow closure. For each biological replicate, the percent of cells that complete furrow ingression was calculated by dividing the number of cells that completed cytokinesis by the total number of cells analyzed. Successful cytokinesis was scored for cells that had at least two stimulated neighbor cells, cells in the same field of view where the neighbors were not included in the region of stimulation (unstimulated neighbors), or cells expressing the optogenetic constructs but in embryos that were never exposed to light stimulation (unstimulated tissue). Cells that completed furrow ingression before stimulation began, or cells that initiated furrowing after stimulation had finished were also included in the unstimulated neighbor group.

### Figure preparation

Images were processed in Fiji. Maximum intensity projections from multiple z-slices are shown. Channels were independently adjusted to highlight relevant features using linear adjustments. Kymographs were generated using a 5 pixel-wide line across the division plane and the multi- kymograph tool in Fiji. LUTs were applied as indicated on the figures. Figures were assembled using Adobe Illustrator.

### Statistical analysis

Mean and Statistical error of the mean (SEM) was calculated using GraphPad Prism. To determine significance, unpaired t-tests were performed using GraphPad Prism or Microsoft Excel.

---

**Supplemental Figure 1:**
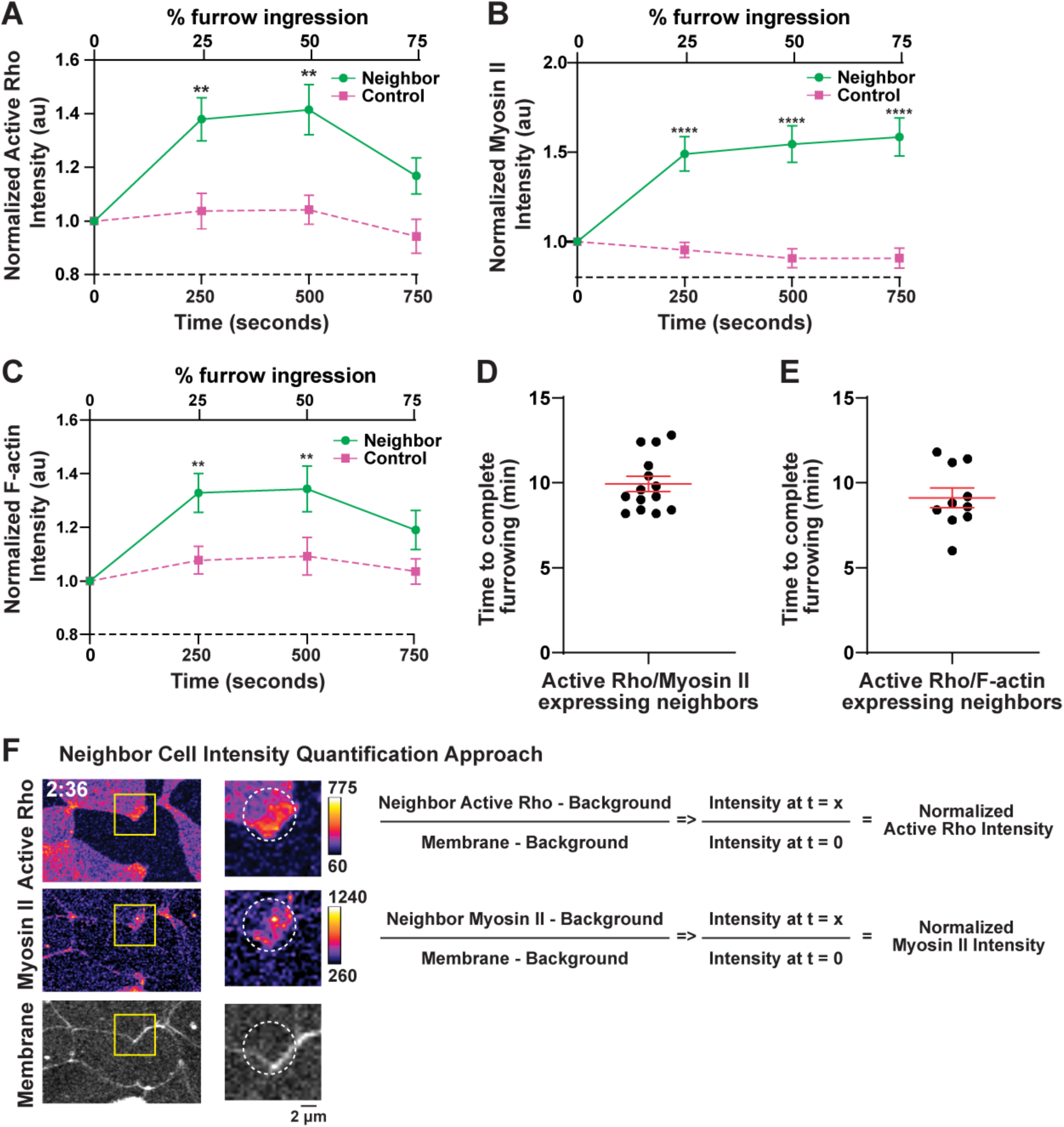
Quantification of neighbor cell intensities during furrowing. A) Quantification of normalized junctional active Rho intensity in neighbor cells at 0%, 25% 50% and 75% ingression (solid x-axis, top). Normalized junctional active Rho intensity in non-dividing control cells measured at 0, 250, 500, and 750 seconds (dashed x-axis, bottom) with the start of furrow ingression as time = 0. For non-dividing controls cells: n = 20 junctions, and for dividing cells: n = 20 junctions from 12 dividing cells across 5 embryos. ** p ≤ 0.01. B) Quantification of normalized junctional Myosin II intensity in neighbor cells at 0%, 25% 50% and 75% ingression (solid x-axis, top). Normalized junctional Myosin II intensity in non-dividing control cells was measured at 0, 250, 500, and 750 seconds (dashed x-axis, bottom) with the start of furrow ingression as time = 0. For non-dividing controls cells: n = 20 junctions, and for dividing cells: n = 20 junctions from 13 dividing cells across 6 embryos. **** p ≤ 0.0001. C) Quantification of normalized junctional F-actin intensity in neighbor cells at 0%, 25% 50% and 75% ingression (solid x-axis, top). Normalized junctional F-actin intensity in non-dividing control cells was measured at 0, 250, 500, and 750 seconds (dashed x-axis, bottom) the start of furrow ingression as time = 0. For non-dividing controls cells: n = 20 junctions, and for dividing cells: n = 20 dividing junctions from 12 cells across 6 embryos. ** p ≤ 0.01. D) Time to complete furrowing for cells used to quantify Active Rho and Myosin II intensities in neighbor cells in Figure 1E. Each dot represents one cell. Mean ± SEM is shown in red. E) Time to complete furrowing for cells used to quantify Active Rho and F-actin intensities in neighbor cells in Figure 1F. Each dot represents one cell. Mean ± SEM is shown in red. F) Approach for quantifying normalized intensities for neighbor cell active Rho and Myosin II (shown in example in F) or active Rho and F-actin (not shown in F) as shown in Figure 1E-F and Figure S1A-C. A 5 µm region of interest (ROI, dashed circle) was drawn in the neighbor cell so that the ROI encompassed signal from both the junction and the near junctional area. The intensities of Active Rho, Myosin II, F-actin and Membrane were measured at each time point during furrow ingression. A background measurement was taken using the same size ROI at a non-expressing or non-stained region of the field of view. The Active Rho, Myosin II, and F-actin intensities were normalized to the membrane signal to account for the changing junction shape at the furrow, and the intensity at each time point was normalized to the start of furrow ingression (t = 0).

**Supplemental Figure 2:**
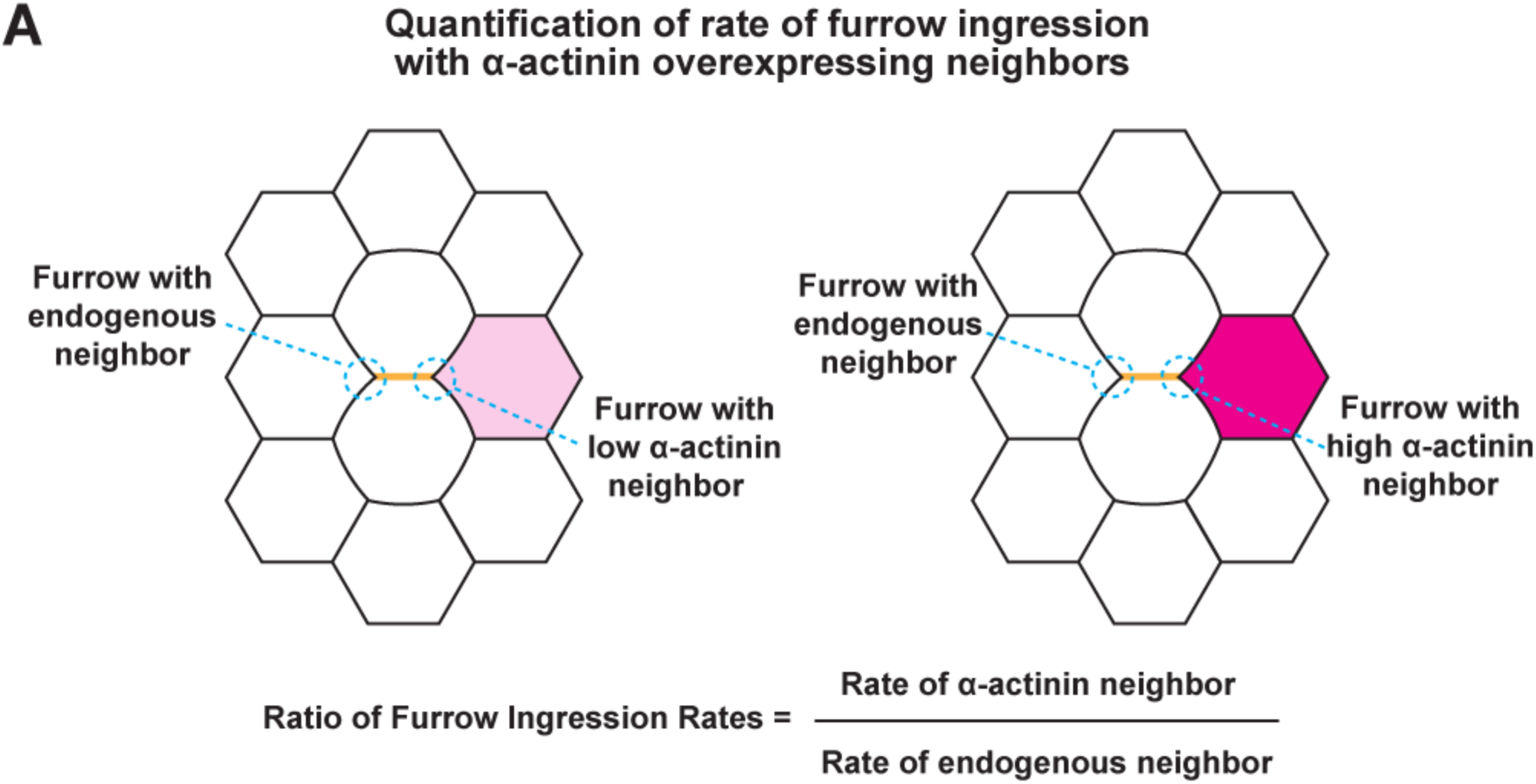
Quantification of rate of furrow ingression when neighbors overexpress ɑ-actinin. A) Top left: schematic of a dividing cell with one neighbor expressing endogenous levels of ɑ-actinin and one neighbor overexpressing low levels of GFP-ɑ-actinin. Top right: schematic of a dividing cell with one neighbor expressing endogenous levels of ɑ-actinin and one neighbor overexpressing high levels of GFP-ɑ-actinin. Bottom: equation used to determine the ratio of furrow ingression rates when a dividing cell has two different neighbors.

**Supplemental Figure 3:**
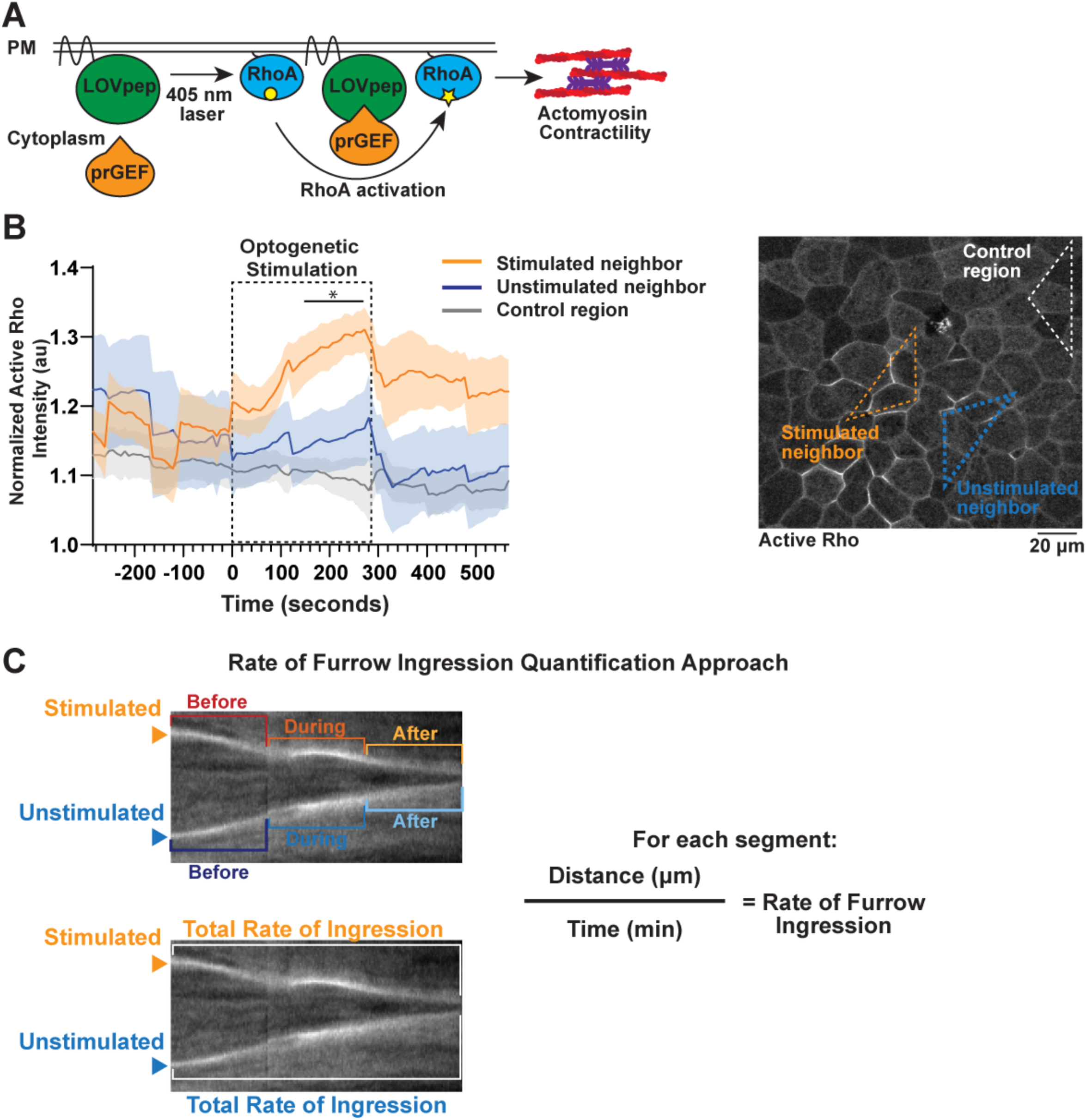
Quantification of response to TULIP optogenetic activation of neighbor cells. A) Schematic of TULIP-dependent optogenetic activation of Rho-mediated actomyosin contractility. B) Left: Plot of active Rho (mCherry-2XrGBD) intensity over time before (-286-0 sec) during (0-286 sec, dashed box) and after (286-564 sec) optogenetic stimulation. Mean intensity (solid line) ± SEM (shading) is shown. active Rho intensity was measured in stimulated neighbor cells (orange), corresponding unstimulated neighbors (blue), and a control ROI (gray) well distanced from the stimulation region and normalized to whole field active Rho intensity. For all: n = 7 regions across 5 embryos, and * p ≤ 0.05, indicates significance of t-test between stimulated and unstimulated sides of the furrow. Right: Example field of view used for quantification of active Rho intensity with regions of interest (control, stimulated neighbor, unstimulated neighbor) shown with dashed outlines. C) Approach for quantifying rate of furrow ingression for cytokinetic cells with optogenetically stimulated neighbors as shown in Figures 3C and 3D. The position of the cytokinetic furrow was tracked over time by kymographs, and the change in distance relative to the change in time was plotted as the rate of furrow ingression. Rate of ingression was measured before, during, and after neighbor cell stimulation (Figure 3C) and for the entirety of cell division (total rate of furrow ingression, Figure 3D).

**Movie S1: Active Rho and Myosin II accumulate near the cytokinetic furrow in neighbor cells.** Relates to Figure 1A. Time-lapse confocal imaging of active Rho (GFP-rGBD, top left), Myosin II (Sf9-mCherry, top right), and membrane (CellMask Deep Red, bottom left) in cells neighboring a dividing cell. Lookup table scales are indicated as shown. Time shown in minutes:seconds. Playback at 10 fps.

**Movie S2: Active Rho and F-actin accumulate near the cytokinetic furrow in neighbor cells.** Relates to Figure 1C. Time-lapse confocal imaging of active Rho (GFP-rGBD, top left), F-actin (LifeAct-RFP, top right), and membrane (CellMask Deep Red, bottom left) in cells neighboring a dividing cell. Lookup table scales are indicated as shown. Time shown in minutes:seconds. Playback at 10fps.

**Movie S3: ɑ-actinin overexpression in neighbor cells slows cytokinetic furrow ingression.** Relates to Figure 2A. Time-lapse confocal imaging of GFP-ɑ-actinin (FIRE lookup table) and cell membrane (CellMask Deep Red, green). Left: dividing cell with one neighbor overexpressing ɑ- actinin at low level; Right: dividing cell with one neighbor overexpressing ɑ-actinin at high level. Lookup table scale is indicated as shown. Time shown in minutes:seconds. Playback at 10fps.

**Movie S4: Optogenetic activation of Rho-mediated contractility in one neighbor cell pauses cytokinetic furrow ingression.** Relates to Figure 3A. Time-lapse confocal imaging of an embryo expressing the TULIP optogenetic system (signal not shown), and a marker for active Rho (mCherry-2XrGBD). The orange triangle indicates the region and duration of light stimulation for activating the TULIP system. Time shown in minutes:seconds. Playback at 10 fps.

**Movie S5: Optogenetic activation contractility in neighbors on both sides of the furrow induces cytokinetic failure.** Relates to Figure 4A. Time-lapse confocal imaging of embryo expressing the TULIP optogenetic system (signal not shown), a marker for active Rho (mCherry- 2XrGBD), and a chromatin marker (mCherry-H2B). The orange triangles indicate the region and duration of light stimulation for activating the TULIP system. Time shown in minutes:seconds. Playback at 10 fps.

## Acknowledgements

We would like to thank the members of the Miller lab for their useful feedback on this manuscript, especially Babli Adhikary and Anna Friedman for their contributions to our thinking on regulation of neighbor cell contractility. We thank Lance Davidson for generously providing GFP-ɑ-actinin and consulting with us about mechanical forces in dividing and neighboring cells, Ed Munro for the Sf9 construct, and Torey Arnold for his efforts cloning Sf9-mCherry. This work was supported by NIH 5R01GM112794-08 to A.L.M., an American Cancer Society Postdoctoral Fellowship to J.L. and NIH 1K99GM147826-01 to J.L.

## Author Contributions

Conceptualization, J.L. and A.L.M., Methodology, all authors, Validation, all authors., Formal Analysis, J.L., E.M. S.C., Investigation, J.L., E.M. S.C., Resources, A.L.M., Writing – Original Draft, J.L. and A.L.M., Writing – Review & Editing, all authors, Visualization, J.L., E.M., S.C., Supervision, A.L.M, Project Administration, A.L.M., Funding Acquisition, J.L. and A.L.M.

